# Genome-wide association study of food liking and food preference patterns in young adults

**DOI:** 10.64898/2026.03.25.714302

**Authors:** Phoebe SC Hui, Jiahui Zhang, Liang-Dar Hwang

**Affiliations:** Institute for Molecular Bioscience, The University of Queensland, St Lucia, QLD 4072, Australia; Monell Chemical Senses Center, Philadelphia, PA 19104, USA

**Keywords:** Food liking, Genome-wide association study, Dietary preferences, Sensory perception, Young adults, ALSPAC

## Abstract

Genetic variation contributes to individual differences in food liking and dietary behaviour. Genome-wide association studies (GWAS) have identified genetic variants associated with these traits, but most evidence comes from middle-aged and older populations. Young adulthood is a key life stage during which long-term dietary habits develop, yet the genetic basis of food liking during this period remains largely unexplored. We conducted GWAS of 97 food-liking traits and two derived principal components (PCs) in 2,784 young adults (aged 25) from the Avon Longitudinal Study of Parents and Children. The PCs captured broader plant-based/seafood (PC1) and meat-based (PC2) food preference patterns. GWAS identified 16 genome-wide significant associations across 14 individual food-liking traits and one food preference pattern (PC1). Cross-trait analyses showed that several variants were associated liking for multiple foods. For example, the lentil-associated variant rs76659918 was associated with PC1 and with liking for avocado, honey, plain yogurt, chilli peppers, aubergines, and black olives, whereas burger- and steak-associated variants were linked to multiple meat-based foods and PC2. Exploratory analyses showed that *TAS2R38* bitter-sensitive alleles were associated with lower liking for Brussels sprouts, with limited evidence for associations with other traits. Comparison with food liking GWAS from the UK Biobank (aged 37–73) showed limited replication, with robust evidence only for the grapefruit juice-associated variant rs2264192. This study identifies genetic variants associated with food liking in young adulthood and suggests that genetic influences operate at both the level of individual foods and broader food preference patterns.

## Introduction

Individual differences in food preferences are complex behavioural traits partly shaped by genetic variation. Twin studies estimated that genetic differences account for a moderate to high (up to 78%) proportion of variation in food preferences in childhood (Breen et al., 2006), adolescence (Smith et al., 2016), and adulthood (Vink et al., 2020), depending on the food category and study population. Evidence for specific genetic determinants of food liking initially came largely from psychophysical and candidate gene studies of chemosensory pathways. In particular, variation in taste and olfactory receptor genes has been linked to differences in taste perception and preferences for specific foods and beverages, including *TAS2R38* and bitter vegetables, *TAS2R31/TAS2R19* and grapefruit, and other taste receptor variants with coffee and alcohol (Diószegi et al., 2019; E. L. Feeney et al., 2021). While findings varied across populations, phenotype definitions, and study designs, these complementary approaches provide biologically interpretable evidence linking chemosensory function to food-related sensory and hedonic responses.

Genome-wide association studies (GWAS) have subsequently broadened this work by testing genetic variation across the genome without restricting analyses to predefined candidate pathways. Some loci map to genes with implicated chemosensory mechanisms, including chemosensory receptor genes such as *OR6A2*-*OR10A2* for liking cilantro (Eriksson et al., 2012) and *TAS2R43-TAS2R46* for liking coffee (May-Wilson et al., 2022). However, GWAS have also identified numerous loci outside established taste and olfactory pathways, with some findings implicating neural and reward-related processes (Pirastu et al., 2016; May-Wilson et al., 2022). Together, these findings illustrate that the genetic architecture of food liking is not restricted to established taste and olfactory pathways and support genome-wide approaches for identifying additional genetic influences on food liking.

However, young adults remain underrepresented in large-scale genomic studies of food liking (Eriksson et al., 2012; Pirastu et al., 2015, 2016; May-Wilson et al., 2022). It remains unclear how consistently associations identified in later adulthood generalise to younger populations, with recent cross-cohort analyses of functional variants in taste and olfactory receptors across showing a mixed pattern (Hwang et al., 2026). Of the 540 SNP-food liking associations identified in the UK Biobank (aged 37–73) and testable in 25-year-old ALSPAC participants, 45 were replicated in ALSPAC, including concordant associations for grapefruit, onion, pear, and apple linking, whereas other strong associations identified in the UK Biobank, such as the olfactory receptor gene *OR4K17* and garlic liking, were not observed in ALSPAC (Hwang et al., 2026). While these differences may reflect statistical power and other cohort characteristics, they raise the possibility that the genetic architecture of food liking may differ across age groups, as food liking and sensory perception can themselves change with learning, exposure and age (E. L. Feeney et al., 2021; Hayes, 2026). Importantly, the cross-cohort analyses were restricted to functional variants in predefined taste and olfactory receptor genes, leaving it unclear whether similar patterns extend across the genome.

To address these questions, we investigated the genetic architecture of food liking in young adulthood using data from the Avon Longitudinal Study of Parents and Children (ALSPAC) (Boyd et al., 2013). Food liking was assessed at age 25, a developmental stage characterised by greater independence in food choice and relatively lower prevalence of chronic disease (Winpenny et al., 2018). We conducted GWAS of 97 food-liking traits and examined whether identified variants demonstrate food-specific versus food dietary pattern-level effects. We further evaluated their associations with findings in the UK Biobank. By focusing on early adulthood, this study aims to provide new insights into the genetic architecture of hedonic food preferences during this understudied stage of life.

## Methods

### ALSPAC sample and genotyping

ALSPAC is a population-based birth cohort established to investigate genetic and environmental influences on health and development (Boyd et al., 2013; Fraser et al., 2013; Northstone et al., 2025). The study initially recruited 14,541 pregnant women residing in the South West of England with expected delivery dates between April 1991 and December 1992, resulting in 14,062 live births and 13,988 children alive at one year. Subsequent recruitment phases increased the total eligible sample to 15,454 pregnancies and 14,901 children alive at one year. Participants have been followed through repeated questionnaires and clinical assessments.

Genomic DNA was extracted from blood samples collected from both mothers and offspring, with mouthwash samples used for DNA extraction in a minority of offspring who did not provide blood. Genotyping of mothers and offspring was performed using the Illumina Human660W-Quad and HumanHap550 platforms, respectively. The present analyses included offspring genotype data only. Imputation was conducted against the 1000 Genomes European reference panel (Phase 1 Version 3) using IMPUTE v2.2.2. Analyses in the present study were restricted to unrelated individuals (genetic relatedness < 0.05; one individual per related pair removed) and variants with a minor allele frequency > 1% and an imputation INFO score > 0.8.

Ethical approval for the study was obtained from the ALSPAC Ethics and Law Committee and the Local Research Ethics Committees. Informed consent for the use of all data collected was obtained from participants following the recommendations of the ALSPAC Ethics and Law Committee at the time. Participants can contact the study team at any time to retrospectively withdraw consent for their data to be used. Study participation is voluntary and during all data collection sweeps, information was provided on the intended use of data.

### ALSPAC Food liking phenotype

Food liking was assessed at age 25 using the Life at 25+ questionnaire administered to offspring in the ALSPAC cohort. Preferences for 97 food items were measured using a 9- point hedonic scale ranging from 1 (“Extremely dislike”) to 9 (“Extremely like”), with 0 indicating “Never tasted.” Questionnaires were distributed online or by post between October 2017 and October 2018. Of 10,001 eligible participants, 4,398 returned completed questionnaires. For genetic analyses, we included unrelated individuals of European ancestry with available genotype data, age and sex information, and valid liking responses (scores 1– 9). Participants selecting “Never tasted” were excluded from analyses for that specific food item.

Study data were collected and managed using REDCap electronic data capture tools hosted at the University of Bristol (Harris et al., 2009). REDCap (Research Electronic Data Capture) is a secure, web-based software platform designed to support data capture for research studies. The ALSPAC website contains a fully searchable data dictionary and variable search tool: http://www.bristol.ac.uk/alspac/researchers/our-data/.

### Food liking and dietary intake in UK Biobank

UK Biobank is a prospective cohort study comprising over 500,000 participants aged 37–73 years (54.4% females) recruited between 2006 and 2010 from 21 centers across England, Wales, and Scotland (Bycroft et al., 2018). Participants completed baseline questionnaires, underwent clinical assessments, and provided biological samples for biomarker and genetic analyses.

Food liking was assessed in 2019 using an online questionnaire administered to participants who had consented to recontact. The questionnaire included 152 items (140 food and beverage items and 12 additional items related to health-related behaviours). Participants rated each item on a 9-point hedonic scale ranging from 1 (“Extremely dislike”) to 9 (“Extremely like”), comparable to the scale used in ALSPAC. Genome-wide association summary statistics for food liking among European-ancestry UK Biobank participants (May-Wilson et al., 2022) were obtained from the NHGRI-EBI GWAS Catalog (Sollis et al., 2023) on 11 December 2022.

As a secondary exploratory analysis, we examined whether food liking associations identified in ALSPAC also extended to corresponding food intake phenotypes available in the UK Biobank. Dietary intake in the UK Biobank was assessed using i) a touchscreen food frequency questionnaire (FFQ) from all participants during their baseline visits to the assessment center and ii) an online 24-hour recall dietary questionnaire (Liu et al., 2011). The 24-hour recall tool was introduced towards the end of the baseline recruitment period, such that 70,046 participants completed it during their baseline visit (instance 0) (Bradbury et al., 2018); it was subsequently administered as an online follow-up over four cycles, though these repeated measures were not used in the present analysis. For each ALSPAC food-liking trait with a genome-wide significant association, the closest-matching UK Biobank intake trait was identified, using either FFQ- or 24-hour recall-derived measures as appropriate. GWAS of these food intake traits among UK Biobank European participants were identified via the OpenGWAS platform (Elsworth et al., 2020) (accessed 2 March 2026), and genetic associations were extracted using the TwoSampleMR R package (v0.6.21) (Hemani et al., 2017, 2018) from GWAS summary statistics produced by the MRC-IEU bulk GWAS pipeline (Mitchell et al., 2019; Elsworth et al., 2020). For 24-hour recall-derived traits, sample sizes of approximately 65,000 appear consistent with restriction to instance 0 participants following genotype quality control.

### Statistical Analysis

Descriptive statistics (mean and standard deviation [SD]) were calculated for each food- liking trait. Responses coded as “0” (“Never tasted”) were excluded from statistical analyses, as only scores 1–9 reflect evaluative hedonic ratings.

Principal component analysis (PCA) and Pearson correlation analysis were performed to explore patterns of food liking. Each food-liking trait was first adjusted for age and sex using linear regression, and residualised values were used for these multivariate analyses. Because participants who selected “Never tasted” for a food could not provide a valid liking score for that item, PCA was performed on participants with valid responses (scores 1–9) for all 97 food liking items using the prcomp() function in R. Pairwise Pearson correlation coefficients among the 97 age- and sex-adjusted food-liking traits and the derived principal components (PCs) were calculated using all available data for each trait pair (pairwise complete observations). Correlation matrices were visualised using the heatmap() function. All analyses were conducted in R version 4.4.0.

GWAS was conducted separately for each of the 97 food-liking traits and for the primary PCs derived from the PCA using PLINK2 (Chang et al., 2015). For individual food-liking traits, only participants with valid hedonic ratings (scores 1–9) for the corresponding trait were included. GWAS were performed using linear regression under an additive genetic model on the original (non-residualised) trait values, with age and sex included as covariates and the first 10 genetic principal components included to account for population stratification. For GWAS of the principal components, age and sex were not included as covariates, as the components were derived from age- and sex-adjusted traits. Genomic inflation factors (λ) were calculated for each trait (range: 0.9989–1.0293), and Q–Q plots were visually inspected to assess residual population stratification.

Independent genome-wide significant loci were identified using clumping with a 500 kb window and r² < 0.05 at a conventional genome-wide significance threshold of *p* < 5 × 10^-8^, based on the 1000 Genomes Phase 3 European reference panel, implemented in FUMA (Watanabe et al., 2017). FUMA was also used to generate Manhattan, Q–Q, and regional association plots, and to annotate genome-wide significant SNPs by identifying the nearest gene and missense variants in linkage disequilibrium (r² > 0.8). The proportion of phenotypic variance explained (R²) by each lead SNP was calculated as 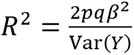, where *p* and *q* are the effect and non-effect allele frequencies, respectively, β is the estimated effect size, and Var(*Y*) is the phenotypic variance of the corresponding trait. Because the food-liking traits were highly correlated, PCA was used to estimate the effective number of independent tests for multiple-testing correction. The first 60 PCs collectively explained more than 90% of the phenotypic variance and were therefore considered to represent the effective number of independent phenotypes. In addition to the conventional genome-wide significance threshold (*p* < 5 × 10^-8^), a Bonferroni-corrected study-wide significance threshold of *p* < 8.3 × 10^-10^ (5 × 10^-8^/60) was also applied.

To characterise the specificity of associations, we examined the effects of lead SNPs across all 97 food-liking traits to determine whether associations were food-specific or extended across multiple food-liking traits, indicating broader food preference patterns. Nominal associations were displayed at *p* < 0.05, *p* < 0.01, and *p* < 0.001 to illustrate the breadth and relative strength of cross-trait associations. Associations surviving Bonferroni correction for 16 lead SNPs and 60 independent tests (*p* **<** 5.2 × 10^-5^; 0.05/(16 × 60)) were additionally highlighted.

As an exploratory analysis, we examined three functional variants in the bitter taste receptor gene *TAS2R38* (rs713598, rs1726866, rs10246939), which influence perception of phenylthiocarbamide and propylthiouracil (Bufe et al., 2005) and have been associated with the perceived bitterness of Brassica vegetables (e.g., Brussels sprouts) (M. A. Sandell & Breslin, 2006) and with a range of food-liking and intake traits (Brito Nunes et al., 2025). Because the three variants were in high linkage disequilibrium (r^2^ > 0.85), they were treated as correlated markers of the same locus. A Bonferroni correction based on the estimated 60 independent phenotypes was applied, corresponding to *p* **<** 8.3 × 10^-4^ (0.05/60).

Genome-wide significant associations identified in ALSPAC young adults were subsequently evaluated in UK Biobank participants of European ancestry for their corresponding food-liking and dietary intake traits using publicly available GWAS summary statistics. Corresponding variants were matched by rsID and harmonised to ensure consistent effect allele orientation. Replication was assessed based on concordant direction of effect and nominal significance (*p* < 0.05). Associations that survived the Bonferroni-corrected threshold for the 16 lead SNPs (*p* < 3.1 × 10^-3^; 0.05/16) were additionally highlighted.

An overview of the study design is presented in **Figure 1**.

**Figure 1.**
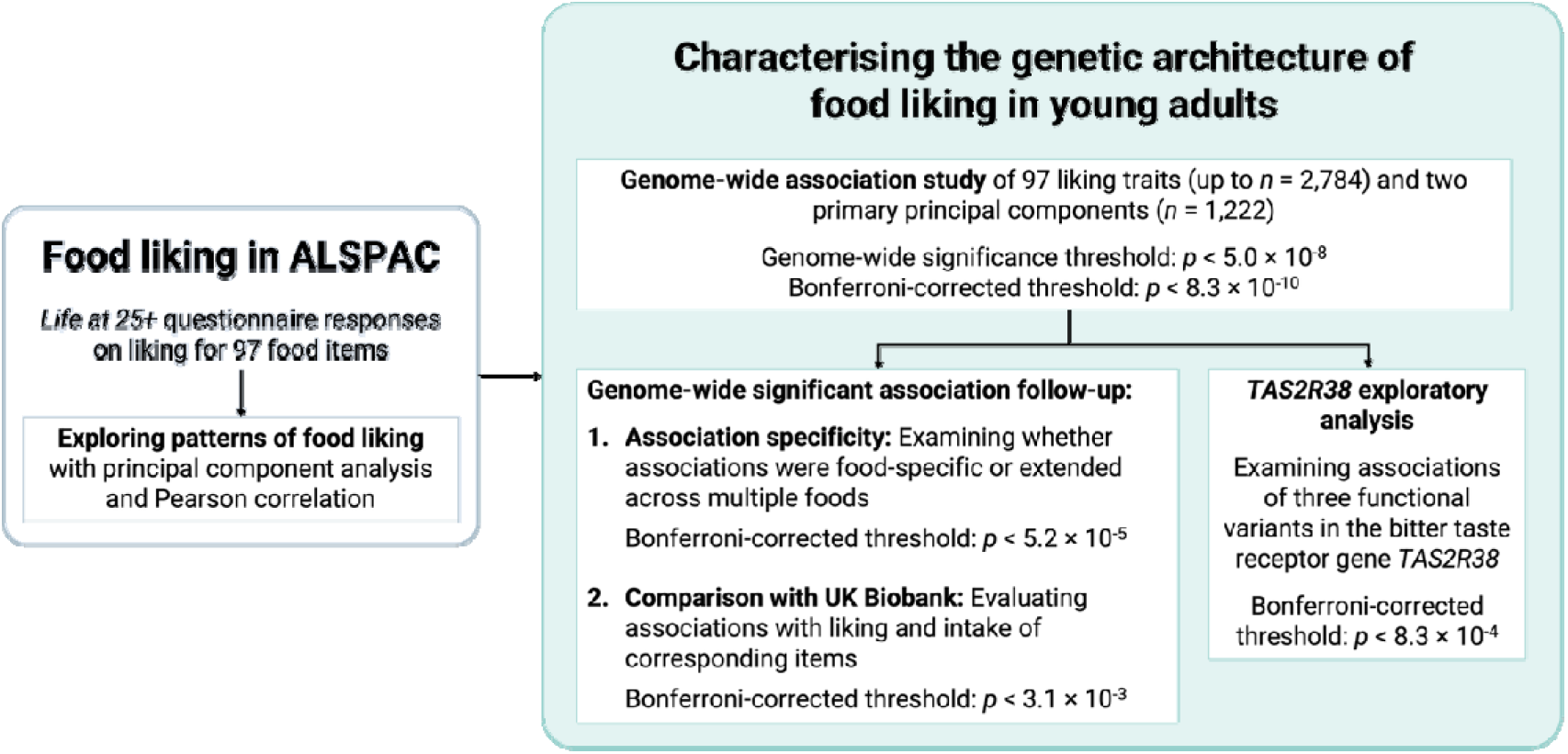
Study design overview.

## Results

### ALSPAC food-liking traits

The final analytic sample included 2,784 participants (mean age = 25.33 years, SD = 0.56; 65.25% females). Mean (SD) food liking scores ranged from 3.40 (2.74) for ale/bitter to 8.00 (1.59) for pizza. The number of participants reporting “Never tasted” varied across items, with the highest frequencies observed for capers (n = 735), artichoke (n = 553), shellfish (n = 330), and high-fiber bar (n = 316). Descriptive statistics for all food-liking traits are provided in **Supplementary Table 1**.

PCA was performed in a subset of 1,222 participants who provided valid ratings for all 97 food-liking traits. The first two PCs (PC1 and PC2) accounted for 21.19% and 8.74% of the total variance, respectively, while each of the remaining PCs accounted for less than 5% (**Supplementary Table 2**). Pearson correlation analyses indicated that PC1 was positively correlated with a diverse set of foods, including liking green olives, black olives, aubergines, avocado, asparagus, smoked fish, shellfish, salmon, prawns, and spinach (r_p_ > 0.60; **Figure 2**; **Supplementary Table 3**), suggesting a broader preference for vegetable, seafood, and other strongly flavoured foods relative to other items in the questionnaire. PC2 was positively correlated with several meat-based foods, including fried chicken, ham, roast chicken, burgers, sausages, bacon, pork, steak, and lamb (r_p_ > 0.60).

**Figure 2.**
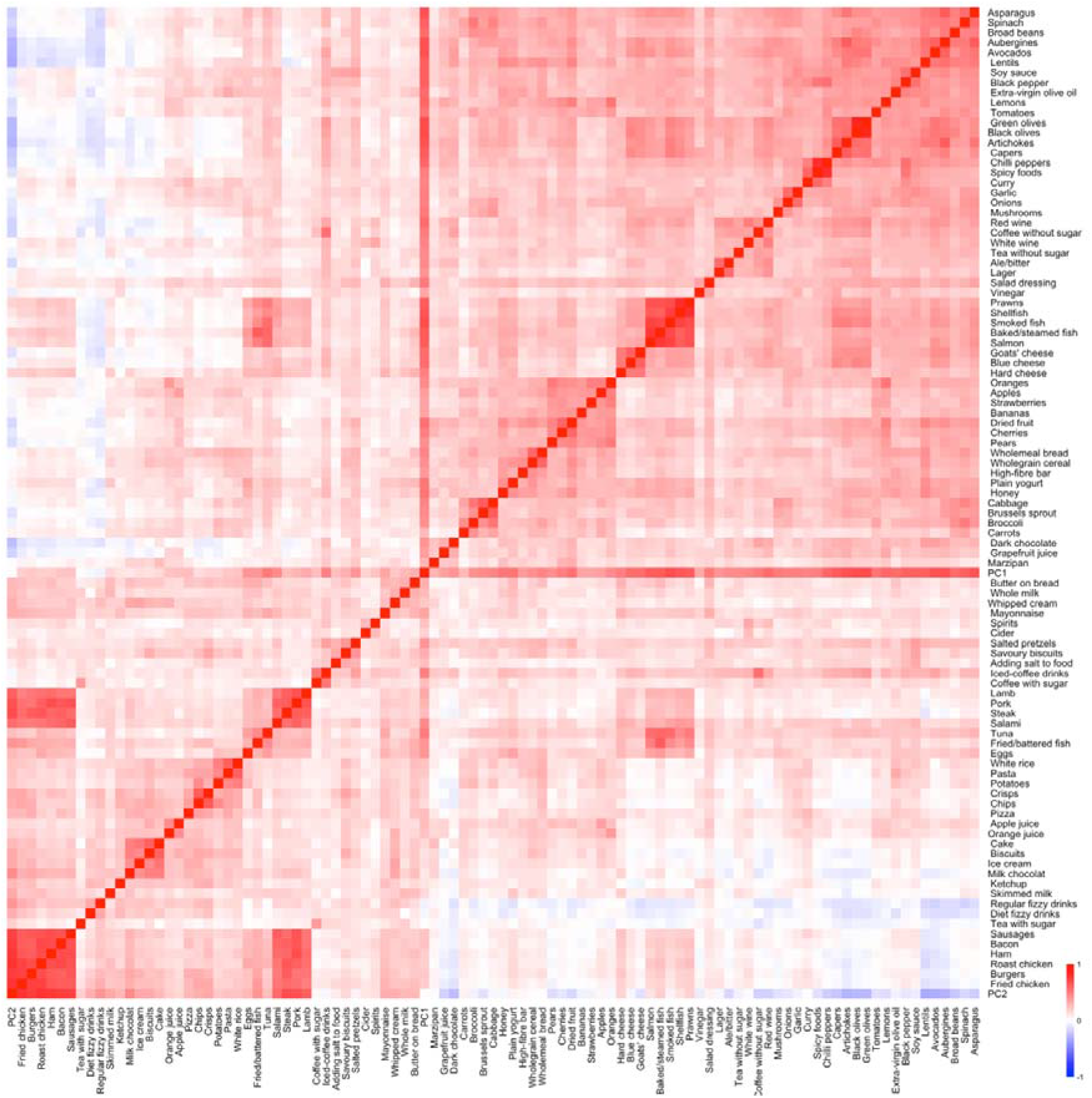
Heatmap showing pairwise Pearson correlation coefficients among 97 food-liking traits in ALSPAC and two food liking principal components (PC1 and PC2). PC1 and PC2 were derived from the principal component analysis of the 97 food-liking traits.

### GWAS of food liking in ALSPAC

GWAS of 97 food-liking traits and two PCs identified 16 genome-wide significant associations (*p* < 5 × 10^-8^; **Table 1**). One association at *PRR4-PRH1-TAS2R14* (rs2264192) for liking grapefruit juice reached the Bonferroni-corrected significance threshold (β = 0.184, Standard Error [SE] = 0.028, *p* = 4.18 × 10^-11^). This variant is in linkage disequilibrium with missense variants within bitter taste receptor genes *TAS2R30* (rs2599404; β = 0.175, SE = 0.028, *p* = 2.28 × 10^-10^), *TAS2R19* (rs10772420; β = 0.173, SE = 0.027, *p* = 3.19 × 10^-10^), and *TAS2R31* (rs10845293; β = 0.173, SE = 0.027, *p* = 2.87 × 10^-10^). Independent SNPs with p < 1 × 10^-6^ from each locus are provided in **Supplementary Table 4**.

**Table 1.** Genetic variants associated with food-liking traits in ALSPAC with *p* < 5 × 10^-8^.

| Food-liking trait | SNP | Chromosome:Position | EA:NEA | EAF | Beta | SE | P | R<sup>2</sup> | INFO | Nearest gene (GENECODE) |
| --- | --- | --- | --- | --- | --- | --- | --- | --- | --- | --- |
| Cider | rs17521099 | 1:226786012 | G:A | 0.080 | -0.312 | 0.051 | 8.94E-10 | 0.20% | 0.966 | *STUM* |
| Grapefruit juice | rs2264192 † | 12:11266342 | C:T | 0.474 | 0.184 | 0.028 | 4.18E-11 | 0.86% | 0.973 | *PRR4, PRH1, TAS2R14* |
| Prawns | rs71561216 | 8:65000845 | G:GA | 0.092 | -0.259 | 0.047 | 4.71E-08 | 0.19% | 0.980 | *RP11-32K4.1* |
| White rice | rs28407649 | 17:8743396 | G:A | 0.014 | -0.673 | 0.119 | 1.68E-08 | 0.03% | 0.922 | *PIK3R6* |
| Pasta | rs146830824 | 1:163000200 | T:C | 0.036 | -0.441 | 0.077 | 1.09E-08 | 0.08% | 0.883 | *RGS4* |
| Lentils | rs76659918 | 4:179482449 | T:A | 0.030 | -0.474 | 0.084 | 1.83E-08 | 0.07% | 0.946 | *SNORD65* |
| Plain yogurt | rs76077978 | 19:35724038 | A:G | 0.017 | -0.618 | 0.109 | 1.43E-08 | 0.04% | 0.911 | *FAM187B* |
| Wholegrain cereal | rs200896019 * | 7:178720 | A: ACAGCAGGCGCT | 0.061 | -0.347 | 0.006 | 4.99E-08 | 0.28% | 0.809 | *FAM20C* |
| Wholemeal bread | rs75162374 * | 16:83282384 | A:G | 0.012 | -0.720 | 0.131 | 4.48E-08 | 0.03% | 0.864 | *CDH13* |
| Crisps | rs149190507 | 7:51578954 | A:G | 0.016 | -0.643 | 0.112 | 1.10E-08 | 0.04% | 0.916 | *COBL* |
| Burgers | rs7193528 | 16:80953912 | G:A | 0.036 | -0.445 | 0.081 | 3.95E-08 | 0.08% | 0.971 | *TSPAN9, RP11-314O13.1* |
| Steak | rs76631755 | 3:145257646 | G:C | 0.119 | -0.234 | 0.043 | 4.82E-08 | 0.24% | 0.945 | *RP11-88H10.2* |
| Steak | rs2943169 | 8:88193740 | A:C | 0.011 | -0.847 | 0.140 | 1.75E-09 | 0.03% | 0.877 | *CNBD1* |
| Black pepper | rs6414978 | 5:70679720 | G:A | 0.289 | 0.185 | 0.033 | 1.75E-08 | 0.24% | 0.811 | *BDP1, RP11-136K7.2* |
| Spinach | rs6858144 * | 4:30167370 | G:A | 0.288 | -0.174 | 0.032 | 4.98E-08 | 0.45% | 0.891 | *AC109351.1* |
| PC1 | rs4301004 | 3:133760908 | C:A | 0.191 | 0.302 | 0.051 | 5.01E-09 | 0.87% | 1.000 | *SLCO2A1* |

Abbreviations: SNP, single nucleotide polymorphism; EA, effect allele; NEA, non-effect allele; SE, standard errors; R<sup>2</sup>, variance explained (%); INFO, imputation score.
†Indicates the association reached Bonferroni-corrected significance threshold of $p < 8.3 \times 10^{-10}$ . \*Indicates that only one SNP within the locus reached the conventional genome-wide significance threshold of $p < 5 \times 10^{-8}$ .

Regional association plots (**Supplementary Figure 1**) show that several association signals were driven by clusters of SNPs in linkage disequilibrium within the same locus, including associations for liking cider, grapefruit juice, prawns, white rice, lentils, burgers, steak, black pepper, spinach, and PC1. Other traits showed more isolated association signals. Manhattan plots and Q-Q plots are presented in **Supplementary Figure 2**.

### Genetic architecture across food-liking traits

To assess whether genetic signals reflected preferences for individual foods or broader food preference patterns, we examined cross-trait associations of lead SNPs across all 97 food-liking traits (**Supplementary Table 5**). Several variants showed associations with liking for multiple foods. For example, the lentil-associated SNP rs76659918 was associated with avocado and PC1 after Bonferroni correction (*p* < 5.2 × 10^-5^) and with salmon, honey, aubergines, black olives, chilli peppers, and plain yogurt at *p* < 0.001 (**Figure 3**). Similarly, SNPs associated with burgers (rs7193528) and steak (rs2943169) showed associations with meat-based foods, including fried chicken, roast chicken, ham, bacon, and lamb, as well as with PC2 (*p* < 0.001).

**Figure 3.**
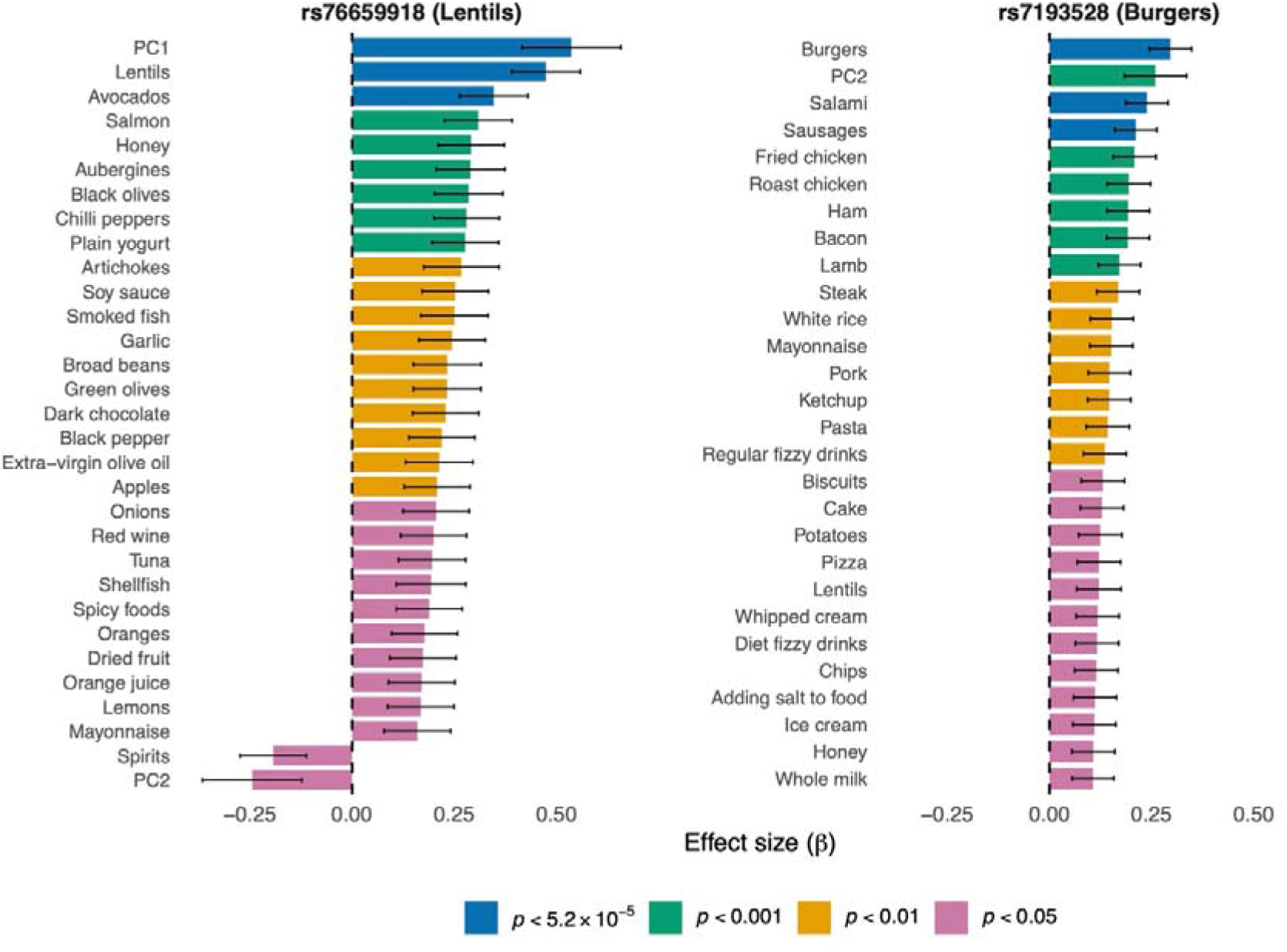
Cross-trait associations for the lentil-associated SNP rs76659918 and the burger-associated SNP rs7193528 across food-liking traits in ALSPAC. Bars represent effect sizes (β) with standard errors. Colours indicate the p-value thresholds (*p* < 2 × 10^-5^, *p* < 0.001, *p* < 0.01, *p* < 0.05). Traits are ordered by effect size within each SNP. Only traits with *p* < 0.05 are shown. PC1 and PC2 were derived from the principal component analysis of the 97 food-liking traits. For visualisation purposes, effect alleles were harmonised so that the lead association for each primary food-liking trait had a positive effect size.

Some loci appeared to have relatively food-specific associations. For example, the white rice-associated SNP (rs28407649) showed no additional associations with other food-liking traits at *p* < 0.01. Similarly, the grapefruit juice-associated SNP (rs2264192) showed a secondary association only with soy sauce (*p* < 0.01), while the cider-associated SNP (rs17521099) showed an association only with spirits (*p* = 0.009).

In the exploratory analysis of *TAS2R38*, the bitter-sensitive alleles (rs713598:G, rs1726866:G, rs10246939:C) were associated with lower liking for Brussels sprouts (*p* < 2 × 10^-5^) (**Supplementary Table 6**). At a nominal threshold (*p* < 0.05), these alleles were also associated with lower liking for spinach and higher liking for biscuits. No evidence of association was observed for liking broccoli or other foods.

### Genetic associations with food liking and intake in UK Biobank

Among the 14 food-liking traits (excluding PC1) with at least one genome-wide significant association in ALSPAC, 13 had corresponding food-liking traits in the UK Biobank and ten had corresponding food intake traits (**Supplementary Table 7**). Only the association between rs2264192 and liking grapefruit juice replicated in the UK Biobank (*p* = 7.73 × 10^-60^). Two variants showed only nominal associations (*p* < 0.05): rs76077978 for liking plain yogurt and rs2943169 for liking steak (**Supplementary Table 8**). The wholegrain cereal-associated variant rs200896019 was unavailable in both the liking and intake summary statistics, while the wholemeal bread-associated variant rs75162374 was unavailable in the intake summary statistics, and no suitable proxies were identified for either variant. None of the remaining variants showed evidence of association with either liking (**Supplementary Table 8**) or intake (**Supplementary Table 9**) traits in the UK Biobank.

## Discussion

In this study, we performed genome-wide association analyses of 97 food-liking traits and two food preference PCs in young adults from the ALSPAC cohort. We identified 16 genome-wide significant loci across 14 individual food traits and one broader food preference pattern (PC1). Most loci have not previously been reported in GWAS of food liking, expanding the catalogue of genetic loci associated with food preferences in young adults. Cross-trait analyses indicated that several variants were associated with liking for multiple foods, suggesting that genetic influences on food liking may operate both at the level of individual foods and across broader food preference patterns. Comparison with results from a large-scale genetic study of food liking in the UK Biobank showed limited replication, with robust evidence only for the locus associated with grapefruit juice liking.

In our cross-trait analyses, several lead variants showed associations with multiple individual foods, particularly the meat-associated loci, whereas others showed relatively restricted association profiles. This pattern is broadly consistent with the extensive cross-trait pleiotropy reported in the larger UK Biobank food liking GWAS, in which most (∼82%) identified loci were associated with more than one food-liking trait (May-Wilson et al., 2022). Variation in association breadth has also been observed among chemosensory receptor variants, ranging from associations with single foods to multiple food-liking traits (Hwang et al., 2026). However, most secondary associations in the present cross-trait analyses were observed at nominal thresholds and should therefore be interpreted as descriptive rather than evidence of established pleiotropic associations.

The limited overlap between the ALSPAC and UK Biobank findings is not unexpected given differences in cohort characteristics. Food liking is not necessarily stable across the life course, but can be shaped by repeated exposure, associative learning, and social and cultural influences on food experience (Eertmans, 2001; Hayes, 2026; Prescott, 2026). Such changes in the phenotypic and environmental context of food liking could contribute to differences in the magnitude of genetic associations observed across life stages. However, age cannot be isolated from other differences between these cohorts. In particular, the ALSPAC and UK Biobank participants represent different birth cohorts and are therefore likely to have experienced different food environments, availability, and cultural norms during development. A separate explanation for the lack of replication is that some ALSPAC associations may not represent robust genetic effects. The relatively modest ALSPAC sample size increases susceptibility to imprecise or inflated discovery effect estimates and false-positive findings. Given the substantially larger UK Biobank sample, the absence of association in the UK Biobank – where the variant and food liking phenotype were sufficiently comparable – is less readily explained by inadequate replication power. Further validation in independent cohorts with comparable food liking measures will be necessary to determine the robustness and generalisability of these associations. Longitudinal analyses within the same birth cohort, or across multiple cohorts with overlapping age ranges, would be needed to determine whether genetic associations with food liking genuinely vary across the life course.

As liking is positively correlated with consumption (May-Wilson et al., 2022), genetic influences on liking may also extend to food intake, although perfect correlations or equivalent magnitude in associations would not necessarily be expected because consumption is additionally shaped by behavioural, social, cultural, and environmental factors (Hayes, 2026). Although lead variants identified from a GWAS cannot be assumed to causally influence the phenotype, examining associations between liking-associated lead variants with intake provides an exploratory means of assessing whether the same genetic signals extend across liking and intake, and if so, the extent of this overlap. However, we observed little evidence that the ALSPAC lead variants were associated with corresponding UK Biobank intake traits. These null findings are difficult to interpret as most of the underlying ALSPAC liking associations did not themselves replicate in the UK Biobank and may not represent robust genetic associations. Accordingly, null findings should not be interpreted as evidence that such carry-over does not occur, but rather the present analysis had limited ability to determine whether genetic influences on liking carry over to food intake.

Despite the limited overlap, we observed evidence of replication for the locus near rs2264192 associated with grapefruit juice liking. This variant is in high linkage disequilibrium with missense variants in the bitter taste receptor genes *TAS2R30* (rs2599404; r^2^=1), *TAS2R19* (rs10772420; r^2^=0.978), and *TAS2R31* (rs10845293; r^2^=0.978). The *TAS2R30* variant has previously been associated with the perceived intensity of quinine (Hwang et al., 2018; Einarsson et al., 2026), with alleles associated with higher perceived quinine intensity also associated with lower liking for grapefruit juice. Likewise, the *TAS2R19* and *TAS2R31* variants have previously been associated with grapefruit bitterness, grapefruit liking, and quinine bitterness (Hayes et al., 2011, 2015). Together, these findings support a role for genetic variation within the bitter taste receptor cluster on chromosome 12 in shaping individual differences in grapefruit preference. Beyond this replicated chromosome 12 bitter taste receptor locus, most genome-wide significant loci identified in this study have not previously been implicated in food liking and were located near genes without established roles in chemosensory perception or dietary behaviour. These loci warrant replication in independent cohorts and functional investigation to establish their biological relevance.

In the exploratory analyses, the *TAS2R38* bitter-sensitive alleles were associated with lower liking for Brussels sprouts in ALSPAC. This finding is biologically plausible as cruciferous vegetables contain glucosinolate-derived bitter compounds, whose perceived bitterness has been associated with *TAS2R38* genotype (M. A. Sandell & Breslin, 2006; E. Feeney, 2011). Although the association of *TAS2R38* variation with PTC/PROP bitterness is well established, studies examining downstream associations with Brassica vegetable preference or intake have produced mixed findings, with both positive and null associations reported across populations (M. Sandell et al., 2014; Diószegi et al., 2019). Consistent with this heterogeneity, a recent UK Biobank study did not detect an association between *TAS2R38* variants and liking for Brussels sprouts despite a substantially larger sample (Brito Nunes et al., 2025). Together, these findings suggest that while the effect of *TAS2R38* variation on PTC/PROP perception is well established, its downstream associations with the liking of complex bitter vegetables may be less consistent and depend on the specific food examined and study populations.

Several limitations should be considered. First, the sample size of the present study is modest relative to large-scale GWAS, which may limit statistical power to detect variants with small effects and increase uncertainty in effect size estimates. Second, replication against the UK Biobank food liking GWAS (May-Wilson et al., 2022) was limited, partly reflecting differences in cohort characteristics and age distributions between studies. Third, food liking was assessed using self-reported ratings of named foods, which may reflect prior exposure, cultural context, and how participants interpret individual food items. In addition, the exploratory UK Biobank intake analyses of food intake were based on a single 24-hour dietary recall, which may not fully reflect habitual intake. Finally, the sample comprises young adults from a single UK population-based cohort. As with many longitudinal cohort studies, participation may be influenced by socio-demographic factors, and both ALSPAC and UK Biobank exhibit some degree of selection bias (Munafò et al., 2018). The extent to which these findings generalise to other age groups or populations remains to be determined. Future studies in larger and more diverse cohorts with harmonised food-liking measures will be important for confirming these associations and clarifying the genetic architecture of food liking.

In summary, this study identified 16 genome-wide significant loci associated with both individual foods and broader food preference patterns in young adults, expanding the current understanding of the genetic architecture of food liking. Some genetic variants were associated with liking for multiple foods, consistent with broader preference patterns, while others showed more food-specific associations. As larger datasets with harmonised measures of food liking become available, integrating genetic and behavioural data may help clarify the biological mechanisms underlying individual differences in food liking and dietary behaviour.

## Declaration of generative AI in scientific writing

AI-assisted technologies were used to aid readability with selected sentences in the manuscript.

## CRediT authorship contribution statement

Phoebe SC Hui: Investigation, Formal analysis, Data curation, Writing – review & editing. Jiahui Zhang: Formal analysis, Writing – review & editing. Liang-Dar Hwang: Conceptualization, Methodology, Investigation, Formal analysis, Data curation, Writing – original draft, Writing – review & editing, Project administration, Supervision, Resources, Funding acquisition.

## Data availability

The informed consent obtained from ALSPAC (Avon Longitudinal Study of Parents and Children) participants does not allow the data to be made available through any third party maintained public repository. Supporting data are available from ALSPAC on request under the approved proposal number, B3544. Full instructions for applying for data access can be found here: http://www.bristol.ac.uk/alspac/researchers/access/. The ALSPAC study website contains details of all available data (http://www.bristol.ac.uk/alspac/researchers/our-data/).

## Acknowledgements

We are extremely grateful to all the families who took part in this study, the midwives for their help in recruiting them, and the whole ALSPAC team, which includes data collection staff, data and administrations staff, technical managers and the technical staff with the Bristol Bioresource Laboratory, based within the University of Bristol. We thank those who contributed to the survey design and data collection of food preferences and dietary intake from UK Biobank.

