## Supplementary Figure for "Genome-wide association study of food liking and food preference patterns in young adults"

**Supplementary Figure 1. Regional association plots created for genetic variants associated with food liking traits ($p < 5 \times 10^{-8}$) in ALSPAC.**

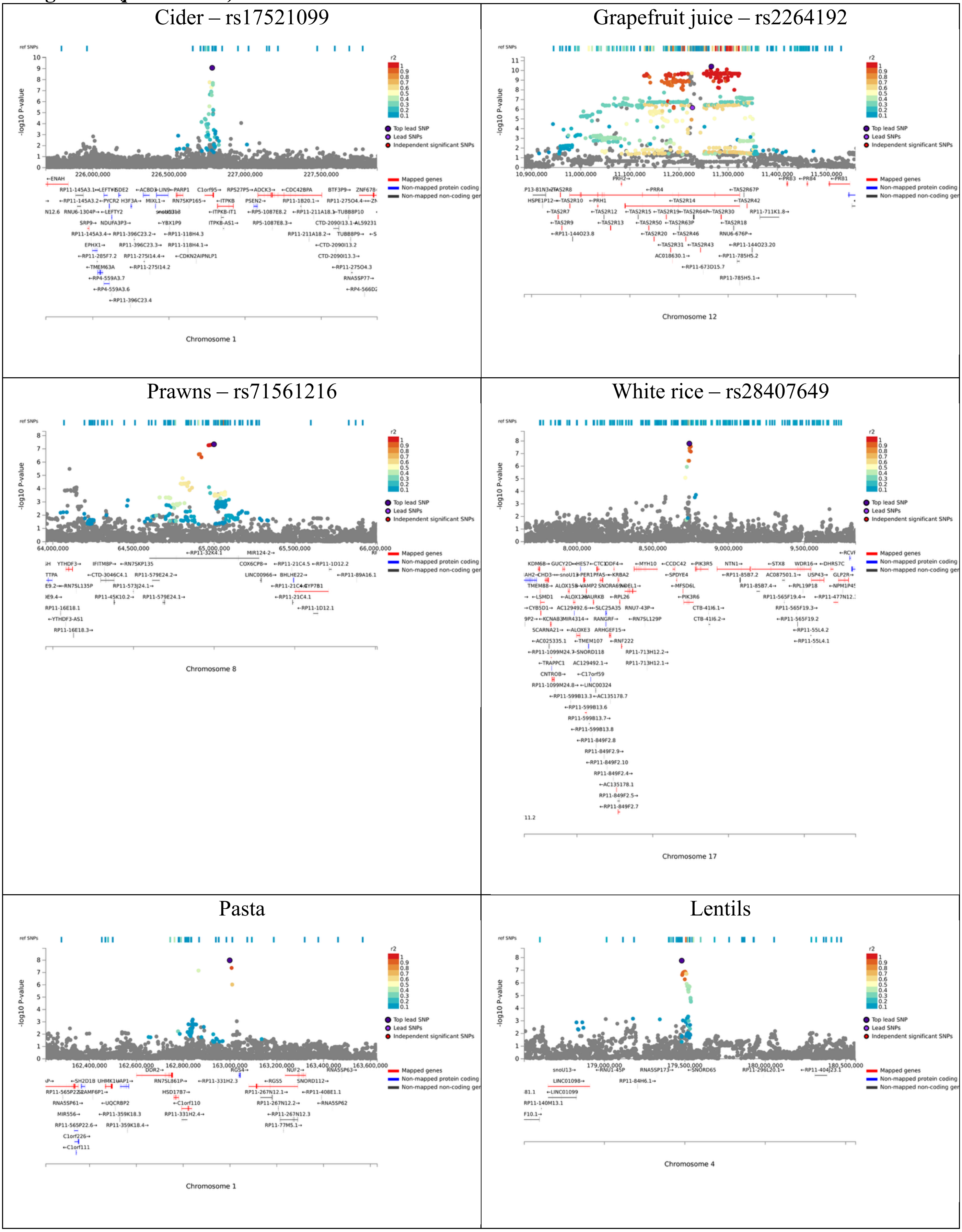

Plain yogurt – rs76077978

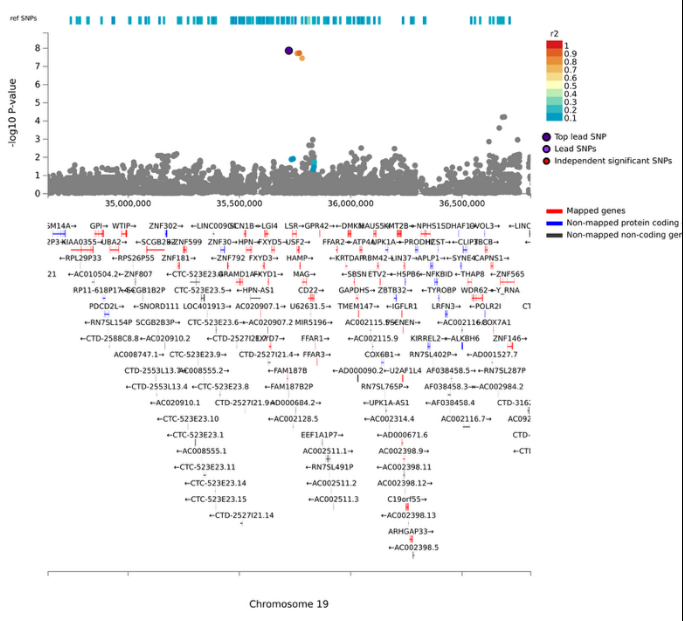


Wholegrain cereal – rs200896019

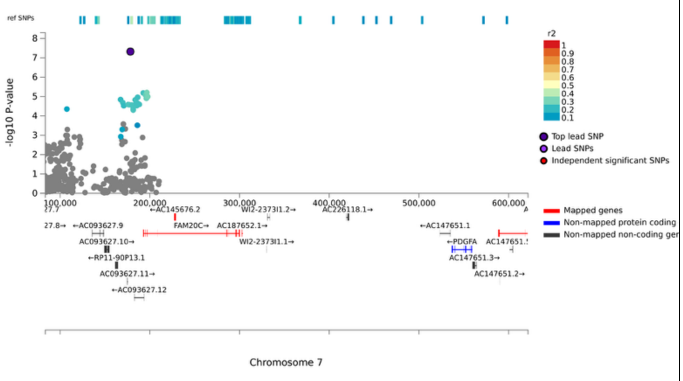


Wholegrain bread – rs75162374

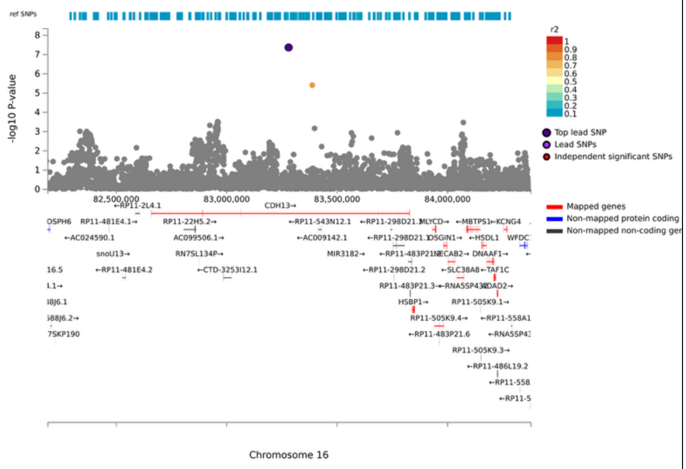


Crisps – rs149190507

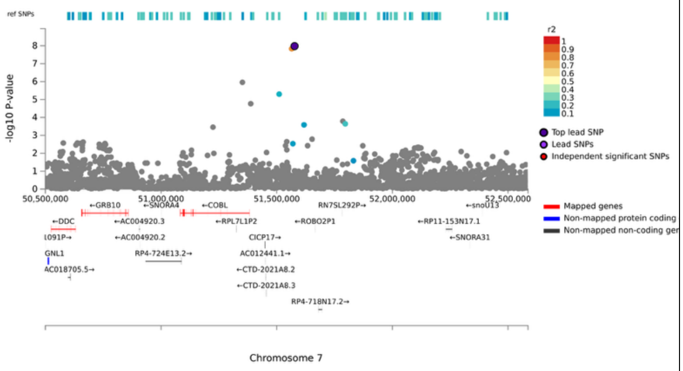


Burgers – rs7193528

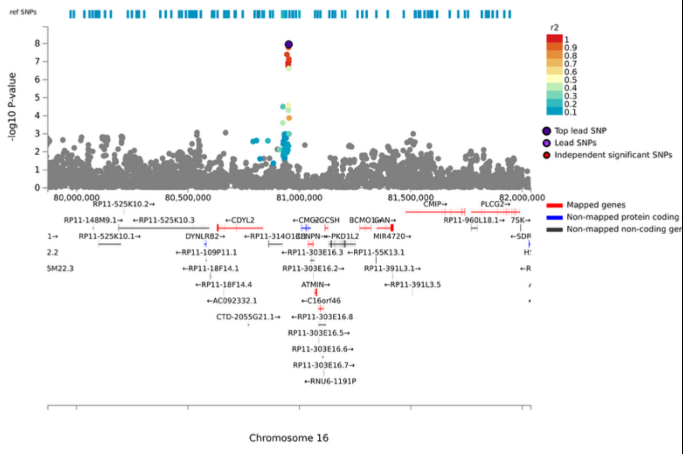


Steak – rs76631755

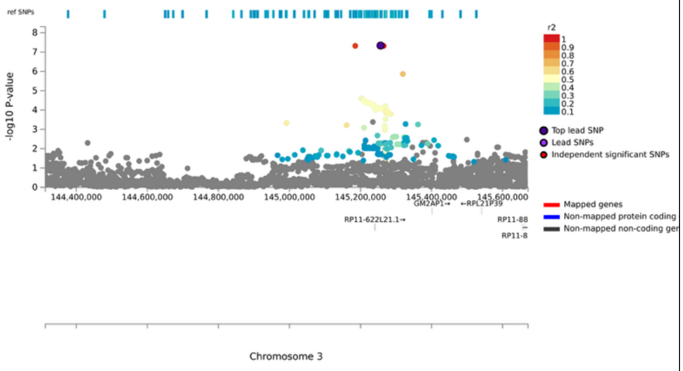

Steak - 2943169

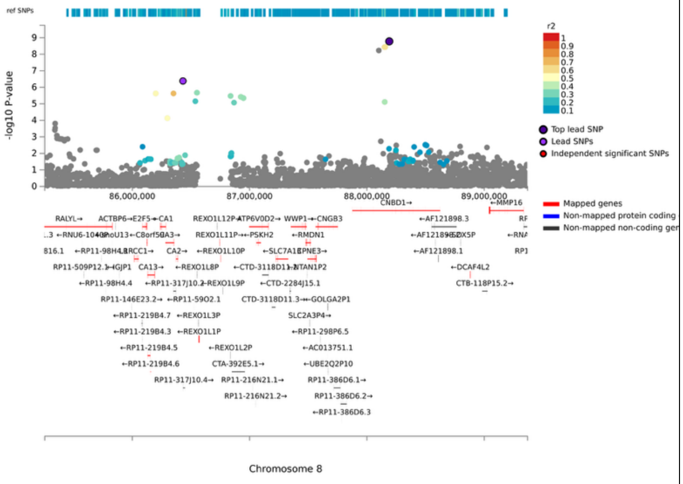


Black pepper – rs6414978

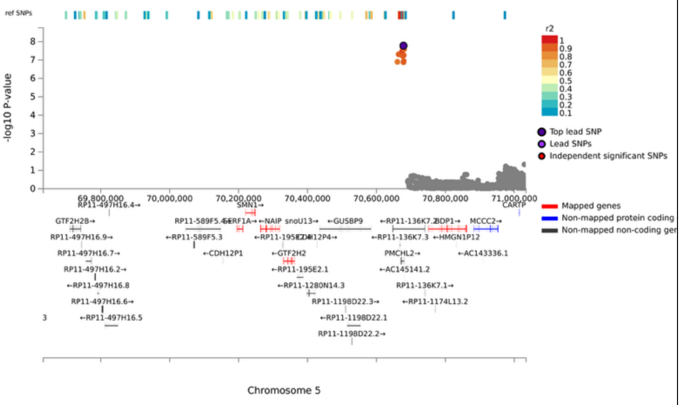


Spinach – rs6858144

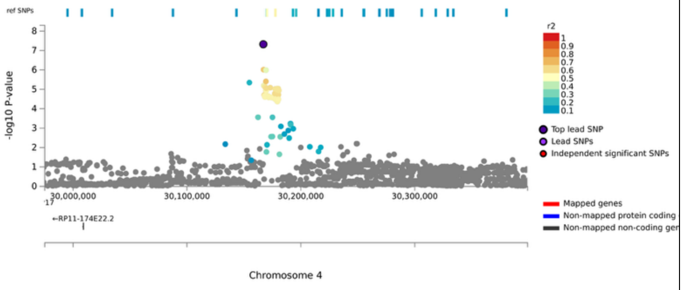


PC1 – rs4301004

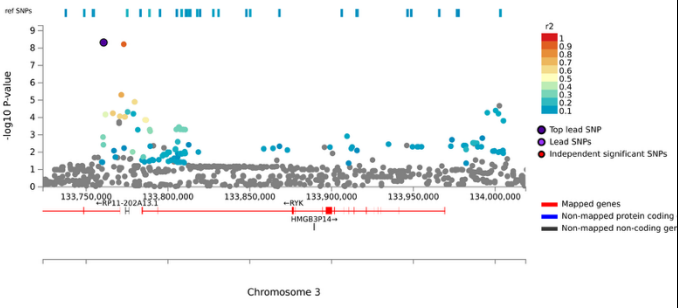

**Supplementary Figure 2. Manhattan and Q-Q plots for food liking traits with at least one genome-wide significant association ($p < 5 \times 10^{-8}$) in ALSPAC.**

Cider

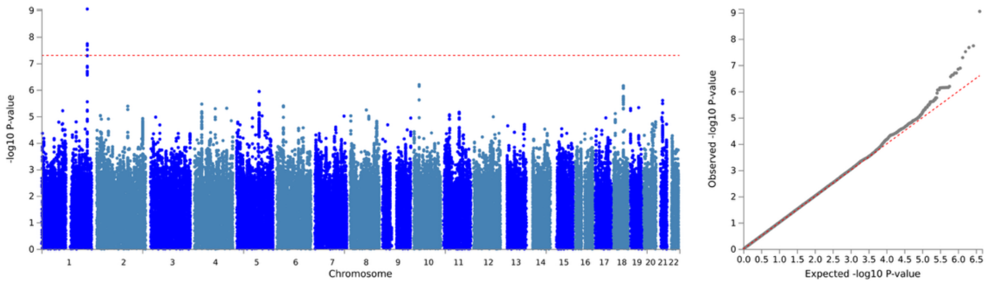


Grapefruit juice

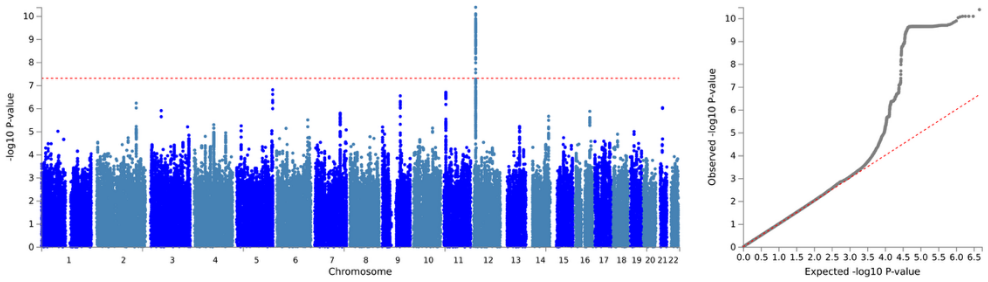


Prawns

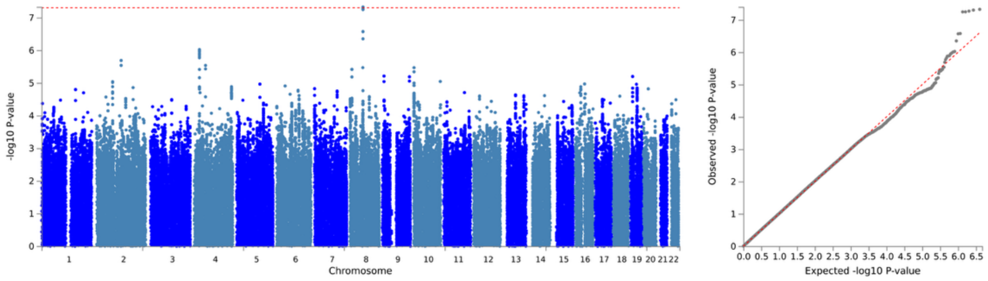


White rice

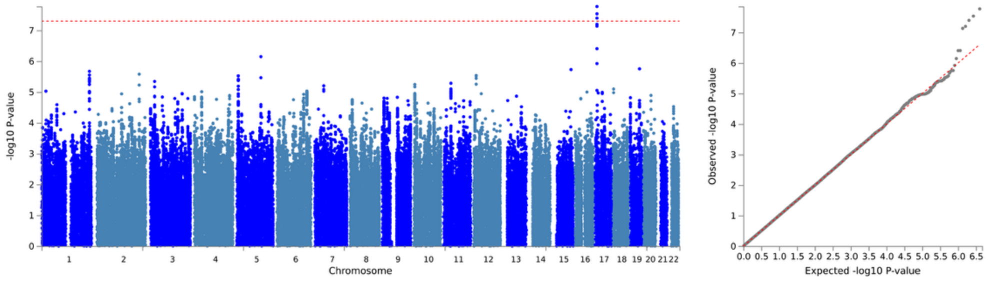


Pasta

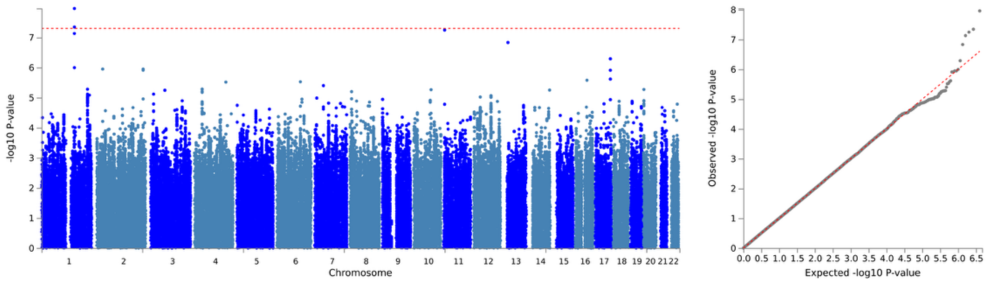

Lentils

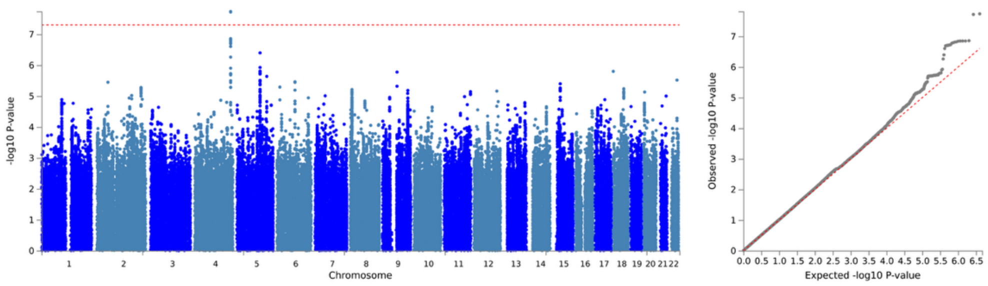


Plain yogurt

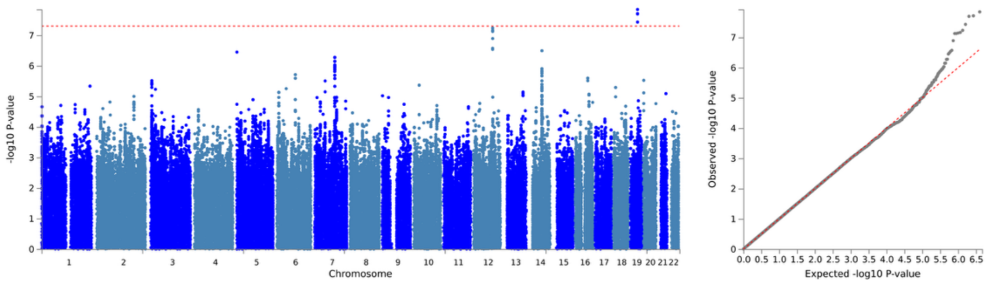


Wholegrain cereal

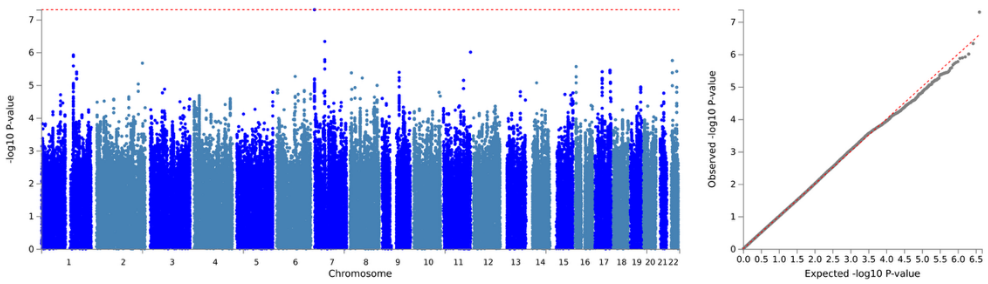


Wholemeal bread

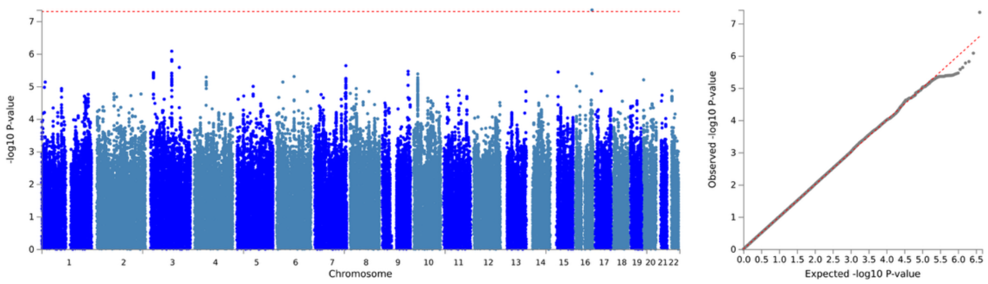


Crisps

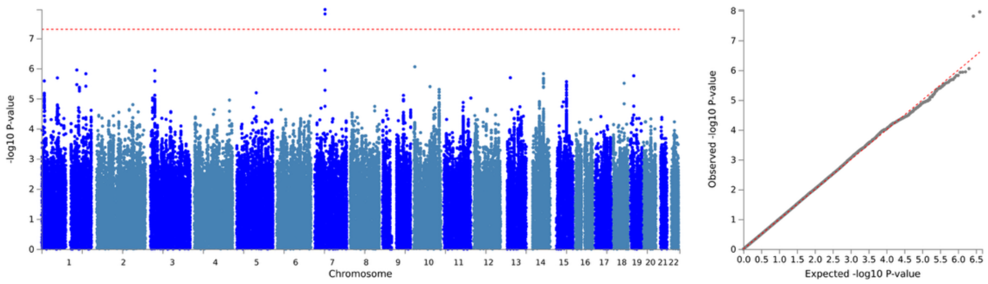


Burgers

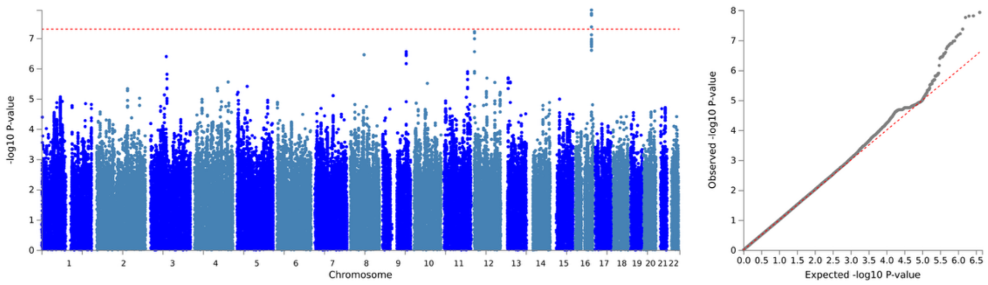

Steak

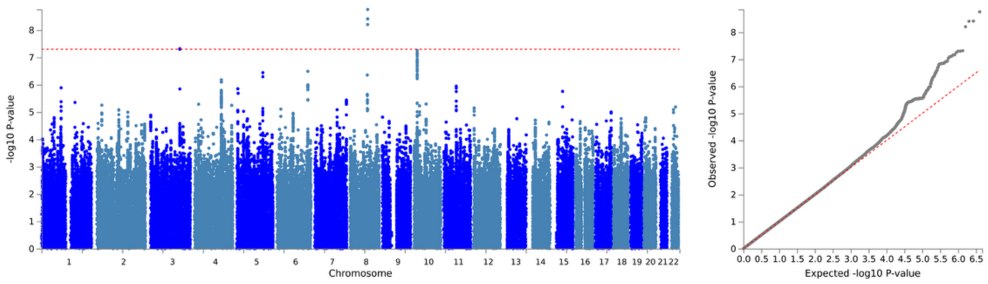


Black pepper

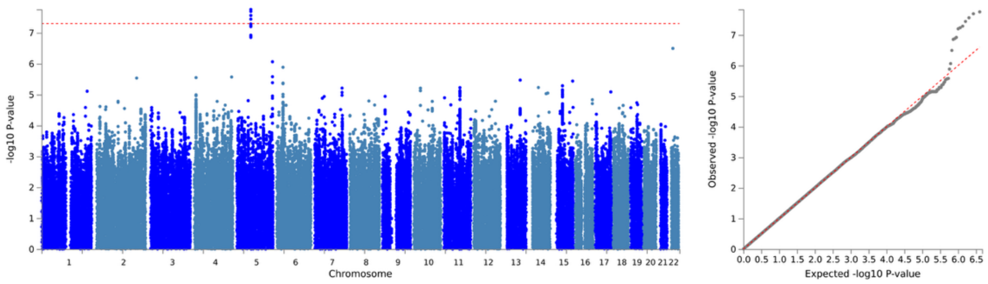


Spinach

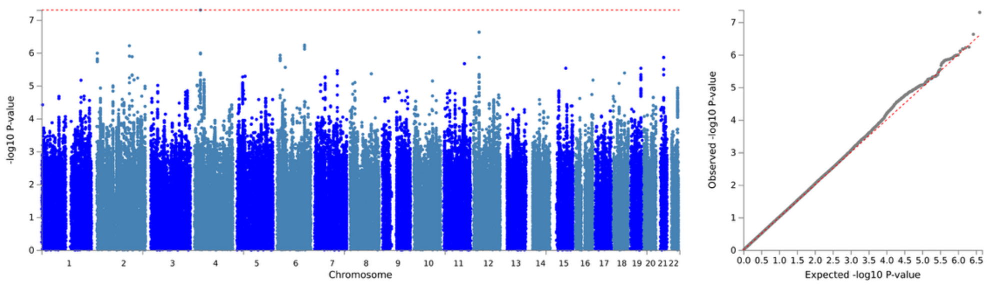


PC1

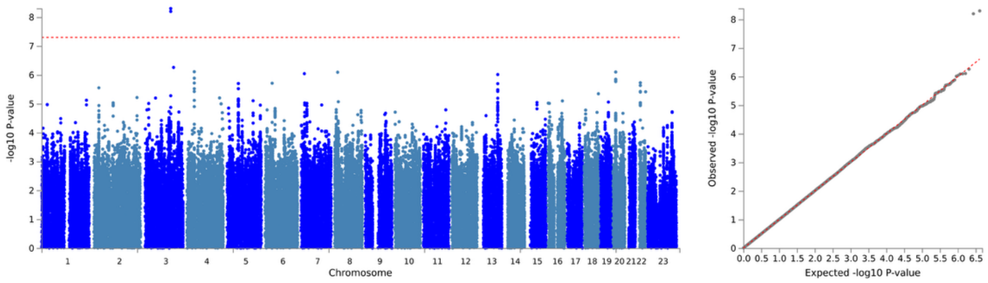
